# The impact of news exposure on collective attention in the United States during the 2016 Zika epidemic

**DOI:** 10.1101/346411

**Authors:** Michele Tizzoni, André Panisson, Daniela Paolotti, Ciro Cattuto

## Abstract

In recent years, many studies have drawn attention to the important role of collective awareness and human behaviour during epidemic outbreaks. A number of modelling efforts have investigated the interaction between the disease transmission dynamics and human behaviour change mediated by news coverage and by information spreading in the population. Yet, given the scarcity of data on public awareness during an epidemic, few studies have relied on empirical data. Here, we use fine-grained, geo-referenced data from three online sources – Wikipedia, the GDELT Project and the Internet Archive – to quantify population-scale information seeking about the 2016 Zika virus epidemic in the U.S., explicitly linking such behavioural signal to epidemiological data. Geolocalized Wikipedia pageview data reveal that visiting patterns of Zika-related pages in Wikipedia were highly synchronized across the United States and largely explained by exposure to national television broadcast. Contrary to the assumption of some theoretical models, news volume and Wikipedia visiting patterns were not significantly correlated with the magnitude or the extent of the epidemic. Attention to Zika, in terms of Zika-related Wikipedia pageviews, was high at the beginning of the outbreak, when public health agencies raised an international alert and triggered media coverage, but subsequently exhibited an activity profile that suggests nonlinear dependencies and memory effects in the relation between information seeking, media pressure, and disease dynamics. This calls for a new and more general modelling framework to describe the interaction between media exposure, public awareness and disease dynamics during epidemic outbreaks.

## Introduction

The advent of the digital era has radically changed the way individuals search for information and this is particularly relevant for health-related information^1^. A 2013 study^2^ found that 59% of U.S. adults had looked for health information on the Web in the previous year and that about one in three U.S. adults use the Internet to figure out what medical condition they have. The fruition of news sources, either traditional such as television, radio and newspapers, or digital such as Web news or online social networks, has become crucial in how health information is delivered and it can play a fundamental role in shaping opinions, awareness and behaviours. In the past ten years, several studies have addressed the impact of awareness and information spread during epidemic outbreaks and it has been reported that the degree of public attention and concern induced by an epidemic threat might affect the disease transmission dynamics ^3–7^. However, modeling efforts have been mostly theoretical and a large-scale empirical characterization of information seeking behaviour and its interplay with the disease dynamics during an epidemic outbreak has been elusive so far due to the lack of available data ^8^.

Here we study a large-scale dataset on spatio-temporally resolved accesses to Wikipedia pages on the 2015-2016 Zika virus (ZIKV) epidemic, regarded as a proxy for collective attention to this emerging health threat. The epidemic started in Brazil in 2015 and spread to other parts of South and North America in 2016. This study focuses on attention patterns in the United States throughout 2016, and on their relation to media coverage of the epidemic.

ZIKV is a RNA virus from the *Flaviviridae* family which is mainly transmitted by infected *Aedes* mosquitoes, although there have been cases of sexual and perinatal transmission. Infection is mostly asymptomatic or associated with mild symptoms ^9^ but it can lead to serious and sometimes fatal neurological defects in neonates born to ZIKV infected women. In particular, following the association between ZIKV and a cluster of microcephaly cases in Brazil^10^, the World Health Organization (WHO) declared the ZIKV epidemic a Public Health Emergency of International Concern (PHEIC) on February 1st, 2016 ^11^. The emergency lasted until November 18th 2016, when the WHO declared the PHEIC to be over ^12^. As of March 2017, ZIKV has spread worldwide to 79 countries where there has been evidence of an ongoing vector-borne virus transmission. The most affected region has been the American continent with 47 countries or territories reporting local ZIKV transmission, due to the extensive presence of *Aedes* mosquitoes in almost all the region’s countries^13^. In such epidemiological context, the ZIKV epidemic has posed peculiar communication challenges to the public due to its association with microcephaly in newborns, its transmission modalities, and its prevalence in areas where the virus was never detected before and that was suddenly characterized by intense international travel due to the 2016 Summer Olympics ^13–15^. Public polls conducted in the United States evidenced the lack of knowledge about ZIKV in the general population and more specifically in groups at risk, such as pregnant women ^16^. The novelty of the disease and the lack of previous knowledge of it in the affected areas make the 2016 ZIKV epidemic an ideal case study to characterize collective attention patterns, identify their drivers and test traditional modeling assumptions. Intuitively, mass media coverage represents the main driver of public attention during an epidemic. Indeed, several peculiarities of media narratives around public health hazards^17^ and infectious diseases^18^ have been elucidated, but a general and quantitative comprehension of how the public opinion responds to media exposure during an emerging epidemic threat is still lacking. The majority of modeling studies assume that media exposure is driving behavioural changes, hence media exposure effects are incorporated into some kind of *media function* that modulates individual behaviors and may affect disease dynamics^19–21^. The general assumption is that as the number of cases increases and is reported by mass media, the susceptibility of individuals will decrease due to increasing awareness and the associated behavioral changes^20^. However, for most disease outbreaks, such an assumption has never been supported by direct empirical evidence. More in general, the complex interplay between media coverage, public attention and disease dynamics during an epidemic remains an open research challenge.

To address these questions, our study analyses time-resolved and geo-localized Wikipedia pageview counts to investigate the dynamics of public attention during the 2016 ZIKV epidemic in the United States. We considered the daily page view counts on 104 different Zika-related Wikipedia articles in the U.S. to be an unambiguous indicator of attention to the epidemic and we investigated the temporal and spatial patterns of page views in relation to the timeline of ZIKV incidence reported by the U.S. Center for Disease Control, and in relation to the coverage of the ZIKV epidemic by local and national media sources. In particular, we focused on news coverage of the ZIKV epidemic by online media and television in 2016, available in digital format through the GDELT project and the Internet Archive (see Methods for a full description of the data under study).

## Results

Public attention and media coverage of the 2016 ZIKV epidemic showed a distinct and synchronous temporal pattern, as seen in Figure 1. The daily timeline of Wikipedia page views (Figure 1 A) highlights two distinct peaks of attention in 2016: the first, in the beginning of February 2016, corresponding to the international alert raised by the WHO and by the CDC at national level; the second, in August 2016, corresponding to the Summer Olympics in Rio de Janeiro, which attracted significant attention due to the health concerns for athletes and the related risk of case importation. Such spikes of attention visibly correspond to similar spikes in the media coverage profiles, both in the TV coverage of the epidemic (Figure 1 B) and in the Web news coverage of Zika (Figure 1 C). The three time series are indeed highly correlated, with a Pearson’s correlation coefficient *r* = 0.74 for the Wikipedia pageviews and the Web news time series, and *r* = 0.80 for the Wikipedia attention and the TV coverage.

**Figure 1:**
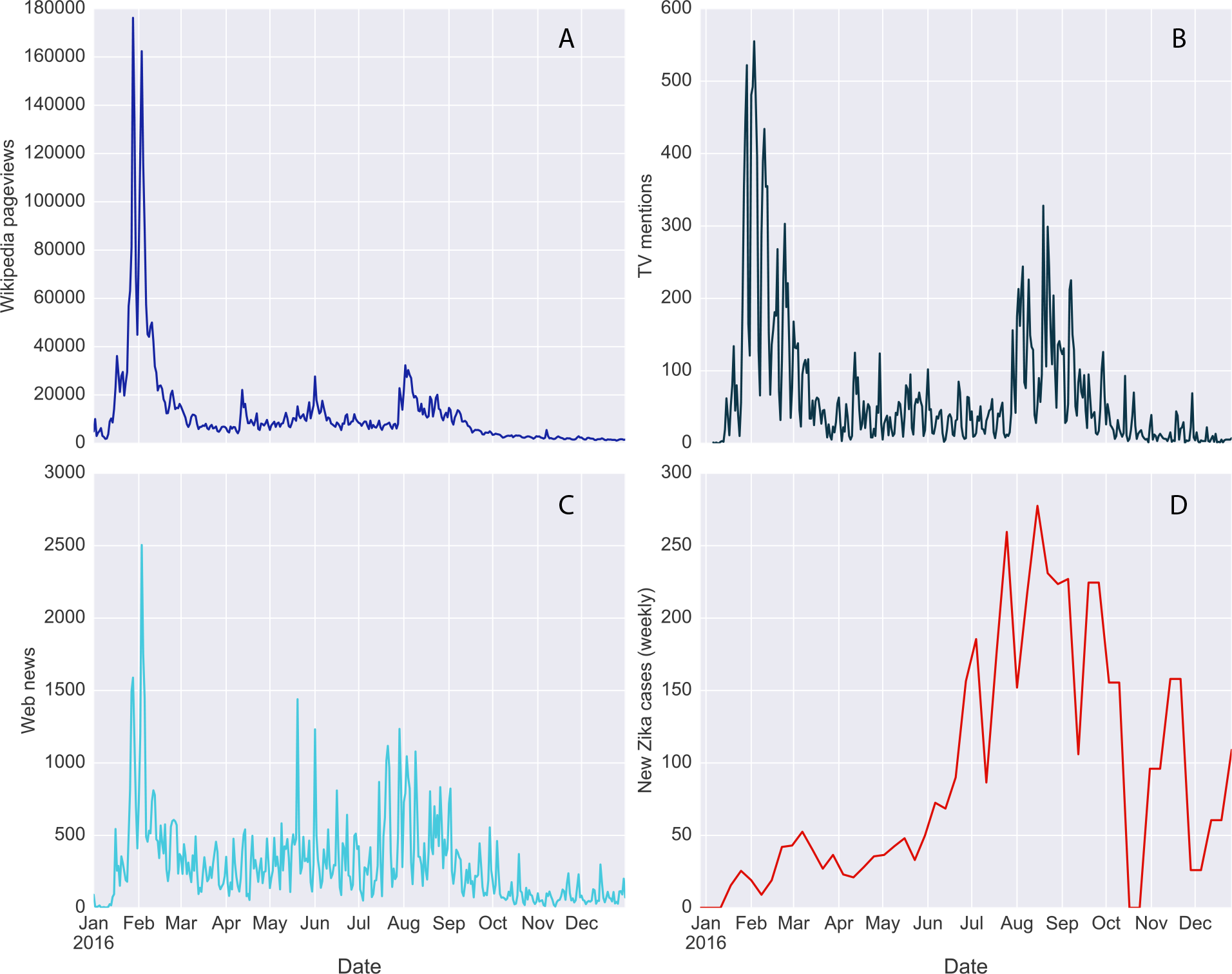
Attention, media coverage and disease incidence of the Zika virus in the USA in 2016. **A,** Daily Wikipedia pageview counts of Zika related pages. **B,** Daily mentions of the word “Zika” in TV programs broadcasted in the U.S. extracted from the TV Internet Archive. **C,** Daily number of Web news mentioning “Zika” extracted from the GDELT project. **D,** Weekly incidence of the Zika virus reported by the Center for Disease Control. Originally reported case counts were smoothed with a biweekly rolling average.

On the contrary, while the profile of Wikipedia pageviews shows a temporal pattern that is very similar to the one displayed by mentions of Zika in media outlets, the temporal profile of the disease incidence is qualitatively very different. The number of new ZIKV cases in the United States reported by the CDC every week (Figure 1 D) gradually increased from the beginning of 2016 until the summer, with a peak between the end of August and the beginning of September. Notifications of new cases declined afterwards. Summer 2016 was also characterized by the first reports of local ZIKV transmission in Florida and in Texas, events that were also responsible for an increased news coverage of Zika. However, the surge of reported ZIKV cases in the United States did not result in an increased level of attention with respect to the initial spike observed at the beginning of the outbreak. The epidemic profile is indeed not correlated with the timeline of Wikipedia page views (*r* = −0.15) or the media coverage profiles (*r* = 0.04 for TV and *r* = 0.14 for Web news). Such dynamics of public attention can be ascribed to the initial novelty of the outbreak, which was presented as a novel and serious health threat even in the presence of a small number of imported cases. As the extension of the outbreak and the associated risks became clearer to the public, the interest of Americans in looking for additional information on Wikipedia faded over the course of the year, with relatively smaller increments linked to important events such as the Olympics. This observation is consistent with the presence of a memory effect in the dynamics of Wikipedia page views: individuals retain information for some time before their attention toward a topic is elicited again by novel events or anniversaries^22, 23^.

The available spatial granularity of Wikipedia page view data allowed us to further inspect how the above picture changes when moving from a national perspective to States and U.S. cities. Notably, the temporal dynamics of attention to the Zika-related Wikipedia pages in 2016 was highly synchronized across all the 50 States. Although the relative risk of case importation and local transmission varied significantly from state to state, being the Southern States more at risk due to vector’s presence and abundance ^24^, the Wikipedia pageview timelines were all highly cor-related, as shown in Figure 2. The Pearson correlation coefficient of the cross-correlation matrix of Wikipedia pageview time series by State ranges from *r* = 0.77 for Delaware and Montana, to *r* = 0.99 for New York and New Jersey. Overall, the correlation of the Wikipedia pageviews in each state with the national timeline was always higher than *r* = 0.88, indicating a high degree of spatial uniformity across the country. Given the above mentioned correlation of Wikipedia pageviews with the TV coverage of the epidemic and the mentions of Zika on the Web, the attention patterns at State level were also highly correlated with the national media coverage suggesting a fundamental role of news exposure as a driver of public attention at all geographic scales.

**Figure 2:**
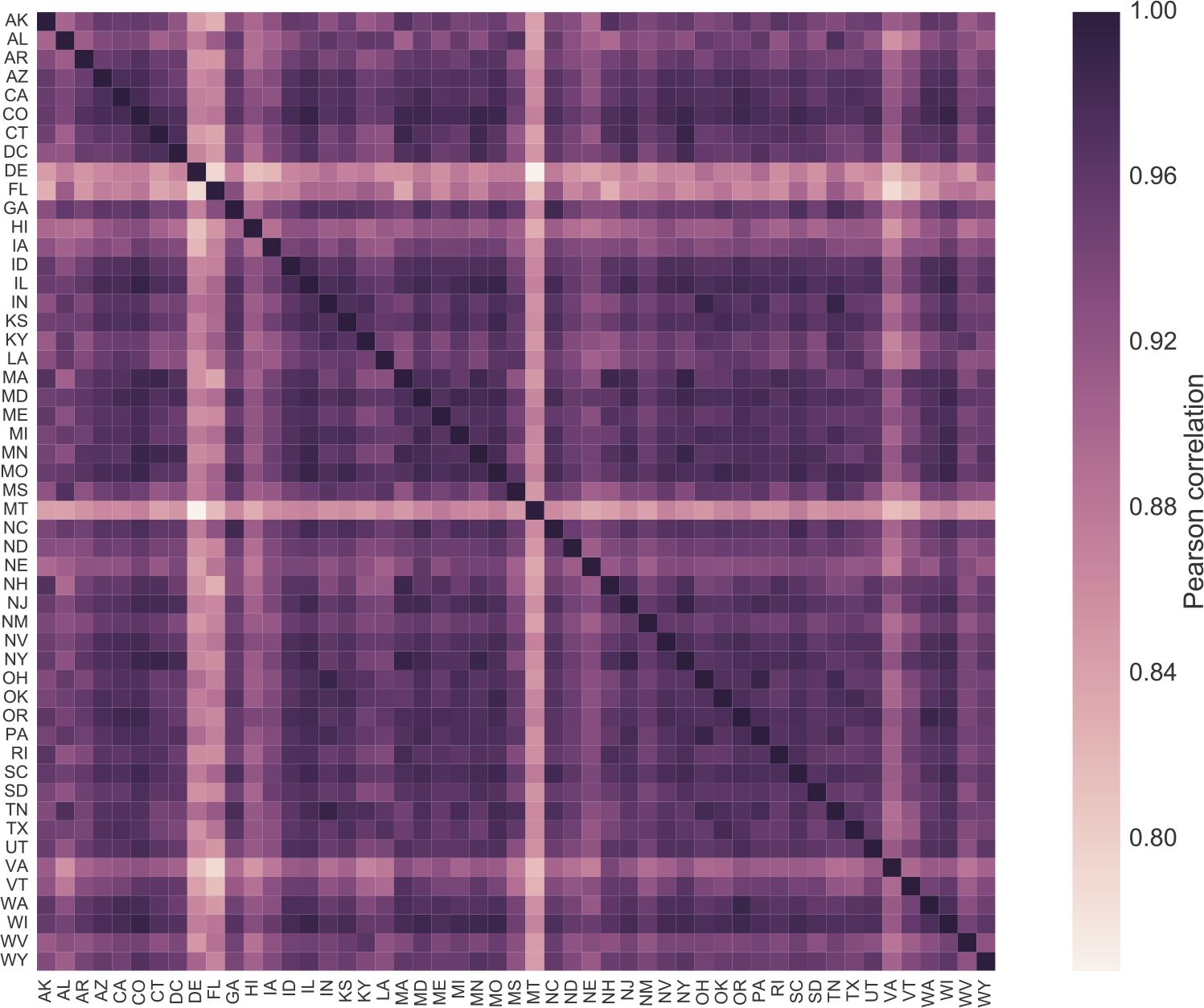
Correlation of public attention timelines by State. Pearson correlation matrix of the daily Wikipedia page views time series of the 50 states and the District of Columbia.

One could argue that local patterns of attention may be influenced by local news and local epidemic events, such as case importations or a local increase of disease prevalence. We tested these hypotheses by comparing Wikipedia page view counts in each state to Web news mentioning the word “Zika” and the name of the state, and to the local ZIKV incidence profiles. Attention profiles in each state were generally positively correlated to Web news mentioning the name of the state, however the degree of correlation ranged from *r* = 0.004 in Wisconsin to *r* = 0.74 in Texas, showing significant spatial differences across the country. Interestingly, ZIKV incidence in each state could explain such geographic variations as a negative driver of attention. On the one hand, local patterns of attention in each state were generally not correlated with disease incidence, with the exception of Montana (*r* = 0.32). On the other hand, Web news covering Zika in each state were positively correlated (*r* > 0.20) with the local incidence profiles only in 20 states out of 50 and, at the same time, these states showed the smallest degree of correlation between news and attention. A direct comparison of the 50 states ranked by degree of correlation between news and ZIKV incidence, and between news and attention, showed a negative rank correlation: weighted Kendall’s *τ* = −0.25. Overall, in those states where local news were following more closely the local epidemic patterns, the dynamics of public attention was not driven much by news. Instead, local attention patterns followed more closely the state news where the latter was more similar to the national one and less correlated with the local ZIKV epidemiology.

It is natural to ask whether correlations between patterns of attention and disease risk may change by looking at different spatial resolutions. To answer this question, we examined the attention to ZIKV in 788 cities of the United States with a population larger than 40,000 and compared it to their total Wikipedia viewership. By ranking the U.S. cities based on their total volume of Wikipedia pageviews in 2016, and comparing such ranking with the one based on pageviews of Zika-related articles only, we identified locations where the attention to ZIKV was higher than expected. As shown in Figure 3 A, cities on the East Coast of Florida showed the highest relative attention to ZIKV, when compared to their overall Wikipedia activity. Other relevant outliers with high attention were cities in Texas and in the Northeast. On the contrary, the lowest attention to ZIKV was observed in cities in California, and in the Midwest (Figure 3B). These results suggest that increases in public attention at city level may be explained by risk perception due to the presence of the vector (as in Florida and Texas). However, the high level of attention in other places, such as Union City, NJ, can not be easily explained by epidemiological risk factors and it may be due to specific events, such as one or more case importations, that do not appear in our dataset.

**Figure 3:**
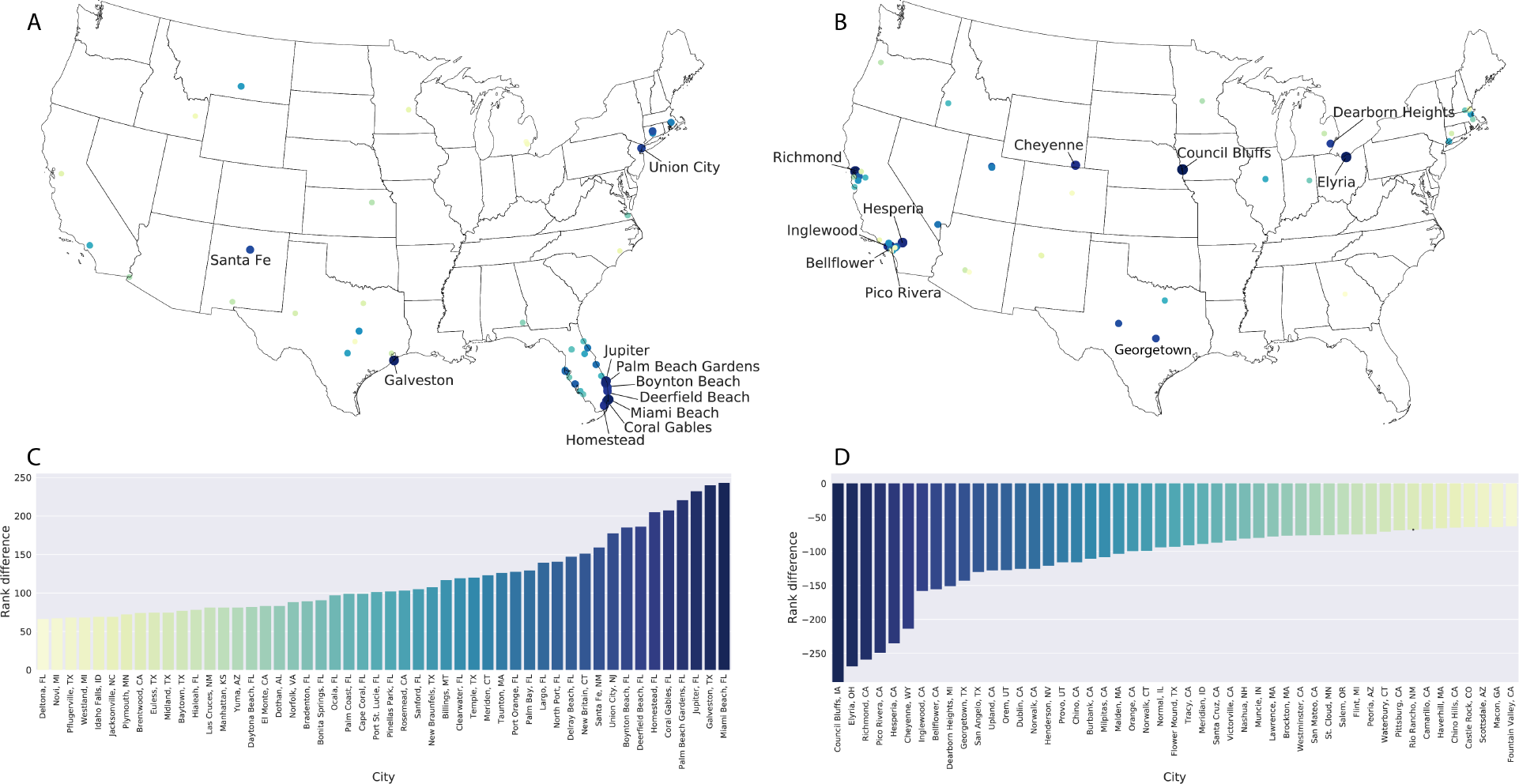
Spatial patterns of attention. Cities of the United States, with population higher than 40,000, where the volume of attention to ZIKV related pages was higher (panel A) or lower (panel B) than expected based on the total volume of page views to Wikipedia in 2016. The maps only show the 50 cities with the largest positive (panel C) or negative (panel D) difference in their page view rankings, based on ZIKV related pages and the full Wikipedia. The labels on the maps highlight the 10 cities with highest (panel A) or lowest attention (panel B).

To gain insight into the relation between media coverage and collective attention, we begin by building an equal-time regression model that predicts the weekly number of Zika-related Wikipedia pageviews for each state, rescaled by state population, based exclusively on the frequencies of Zika-related mentions in Web news and TV closed captions. That is, we assume that information seeking behavior in Wikipedia is driven, at any given point in time, by same-week exposure to media sources. Since our goal is uncovering drivers of collective attention, rather than achieving optimal prediction of the empirical time series, we choose an equal-time modeling approach over standard time series modeling techniques (e.g., autoregressive models). More specifically, we start with a linear regression model that predicts population-rescaled pageview counts for a given week and a given state using only national Web news and TV data for the same week. We focus on 43 states with population in excess of 1 million, comprising more than 98% of the U.S. population according to 2016 United States Census Bureau estimates^25^. We train the model via state-wise cross-validation and evaluate its performance using the determination coefficient *R*^2^ and the Pearson’s correlation coefficient r. Despite its simplicity, this equal-time linear regression demonstrates that both media signals, taken independently, are already quite informative of the Zika population-rescaled pageview time series: using exclusively TV close captions we obtain *R*^2^ = 0.61 and *r* = 0.80, while using only Web news we obtain *R*^2^ = 0.52 and *r* = 0.78. Combining both features, the linear model achieves *R*^2^ = 0.63 and *r* = 0.82. As model performance is evaluated via state-wise cross-validation, these results highlight that national-level media signals are highly informative of state-level pageview time series, once they are rescaled to take into account population size.

To take into account the possibility of memory effects in the response to media exposure, we enrich the feature space of the regression model with additional features (time series) obtained by filtering the Web news and TV time series with an exponential memory kernel (see Methods for a complete description). The characteristic time *τ* of the memory kernel, describing news persistence in the attention response, is a new hyper-parameter of the model to be set via cross-validation. Table 1 summarizes the performance of the model, in terms of determination coefficient *R*^2^ and Pearson’s correlation coefficient *r*, for 10 different sets of features. The introduction of a memory kernel increases the determination coefficient by about 20%, reaching an average *R*^2^ = 0.76. This is obtained for a characteristic time scale *τ* of about 2 weeks, over which collective attention is affected by past media exposure.

**Table 1:** Comparison of model performance with 10 different feature combinations. For each feature, the average *R*^2^, Pearson *r* and AIC, computed over 43 states are reported. Values of *R*^2^ and *r* are computed under K-fold cross-validation (*k* = 10).

| Features | $R^2$ | Pearson $r$ | AIC | $\Delta_i$ |
| --- | --- | --- | --- | --- |
| TV | 0.6091 (0.040) | 0.8011 (0.021) | -1001.90 | 46.67 |
| Web | 0.5244 (0.046) | 0.7780 (0.026) | -986.01 | 62.54 |
| TV, Web | 0.6266 (0.044) | 0.8209 (0.023) | -1003.92 | 44.65 |
| TV, m(TV) | 0.7062 (0.047) | 0.8675 (0.025) | -1025.90 | 22.66 |
| Web, m(Web) | 0.6352 (0.051) | 0.8178 (0.027) | -1004.53 | 44.03 |
| TV, Web, m(TV) | 0.7318 (0.052) | 0.8761 (0.026) | -1033.91 | 14.64 |
| TV, Web, m(Web) | 0.7603 (0.050) | 0.8942 (0.026) | -1045.90 | 2.66 |
| TV, Web, m(TV), m(Web) | 0.7602 (0.050) | 0.8942 (0.026) | -1044.64 | 3.92 |
| TV, Web, m(TV), m(Web), state_news | 0.7638 (0.045) | 0.8937 (0.024) | -1048.56 | 0.0 |

We also considered state-level news features obtained by counting the weekly number of mentions of each state in Web news. However, adding these features does not significantly improve the model predictions (Table 1, bottom row), although it yields the best performance according to the Akaike Information Criterion (AIC). Overall, by computing the AIC for each model and averaging over all states, three linear models based on TV, Web news, and state news, can be considered equally likely, assuming evidence for Δ_*i*_ = *AIC*_*i*_ - *AIC*_*min*_ < 4.

## Discussion

Our study demonstrates that the temporal dynamics of Wikipedia pageviews in the United States during the ZIKV 2016 epidemic was highly predictable, even at state level, based on the volume of national and international news sources mentioning Zika and the United States. Collective attention to the ZIKV outbreak thus seems to have been mainly driven by news exposure and much less by the disease transmission dynamics, although the epidemic profile of ZIKV infections varied significantly from state to state and the risk of local transmission was not uniform across the country. Such picture describes a scenario where the awareness of the epidemic in the country is globally present, while local effects, as those due to the local spreading of awareness, play a less important role ^26^.

Media outlets in the U.S. have a prominent role in defining the on-line public discourse ^27^. The impact of media exposure on the collective awareness and risk perception during epidemic outbreaks has been investigated in previous works ^17,18,28^, however, only a few studies have attempted to quantitatively measure the effect of media engagement on epidemic awareness using empirical data from Web sources on a large scale ^21,29,30^. While previous studies have focused on newspaper coverage of epidemics^31^, we investigated the relationship between the exposure to TV coverage and online news, and the attention to Wikipedia pages. Our study confirms the high sensitivity of Wikipedia searches to breaking news and official announcements, in particular in the case of disaster events, as found by previous studies^32, 33^. On the other hand, the temporal dynamics of Wikipedia page views during the 2016 ZIKV epidemic showed a nonlinear dependence with media coverage: the Wikipedia pages activity was high in the initial phases of the outbreak, but it declined more quickly than media coverage. This can be explained by the fact that information on Wikipedia is rather static, and users will view Wikipedia pages immediately after the news breaks but they will not return in the next days, unless more recent events renew their attention ^32^.

From an epidemiological standpoint, our results are consistent with the recent findings of Bragazzi et al.^34^, who analyzed various data streams to measure the global reaction to the 2015-2016 ZIKV outbreaks in different countries. Similarly, we did not find any statistically significant correlation between the attention to Wikipedia pages and the ZIKV incidence data in the U.S. The correlation between ZIKV incidence and media coverage was also mild, and varied from state to state, suggesting that media coverage was only relatively influenced by the actual progression of the epidemic over time. One might argue that our results may not generalize to all epidemic outbreaks. Indeed, the peculiar characteristics of the ZIKV infection, such as its association to mild symptoms and the relatively small size of the population at risk, due to the spatial distribution of the vector, may have influenced the attention dynamics during the outbreak. Epidemic outbreaks caused by different pathogens, possibly characterized by a higher transmissibility, and different symptomatology, such as the Ebola virus or pandemic influenza, may lead to different attention patterns. However, it is reasonable to believe that media coverage would be, in any case, the main driver of collective attention, as it also has been during the 2014 West African Ebola virus epidemic ^29, 35^.

The increasing availability of novel data streams, such as social media, Web search queries and participatory surveillance data, provides an invaluable resource to measure and quantify the complex interplay between the spread of information, collective attention and the epidemiology of infectious diseases ^36, 37^. Recently, Wikipedia pageview data have been increasingly used by researchers in epidemiology and infectious disease modeling ^38, 39^. The overall value of Wikipedia data to measure and forecast the dynamics of infectious diseases has been debated ^40^ and, in gen-eral, Wikipedia-based forecasting models have been proved successful in the case of endemic or seasonal diseases, such as influenza, dengue or tubercolosis ^39^. On the other hand, our study demonstrates that Wikipedia page viewership can provide a temporally resolved measure of collective attention during epidemic outbreaks caused by novel emerging diseases, at a high spatial granularity. Previous works have investigated the effects of external events on the activity of Wikipedia editors and on the number of pageviews ^41, 42^. More generally, the characterization of the usage of Wikipedia as a source of information and as a proxy for measuring the global attention to real-world events has been studied ^22,33,43,44^. The results of our study add further evidence of the value of Wikipedia data in the field of digital epidemiology, especially for capturing information seeking behavior, and attention patterns during disease outbreaks ^45^.

We showed Wikipedia data can capture collective attention during outbreaks, however, we did not link such signal with a measure of behavioral response in the population. Detecting behavioral changes from Web sources remains a challenging task. Previous studies have used TV viewing data to infer the behavioral response during the 2009 A/H1N1 pandemic in Mexico ^46^. More recently, Poletto el al. ^30^ showed that an increased collective attention was correlated to changes in the hospital management of MERS-Cov patients, reducing the time from admission to isolation. Further research is needed to infer causal patterns between collective attention and behavioral responses, and to identify the most suitable approach to integrate them into disease-behavior models.

## Methods

### Data sources

#### Wikipedia page view counts

We collected hourly pageview data of the English Wikipedia pages “Zika virus” (https://en.wikipedia.org/wiki/Zika_virus) and “Zika fever” (https://en.wikipedia.org/wiki/Zika_fever) and their counterparts in 96 different Wikipedia projects. The complete list of the 104 monitored Wikipedia pages is provided in the Supplementary Information file. While aggregate hourly and daily pageview data for Wikipedia articles by language is released by the Wikimedia Foundation in the form of data dumps (https://dumps.wikimedia.org/other/pageviews/readme.html) and APIs (https://wikitech.wikimedia.org/wiki/Analytics/AQS/Pageviews), the geographic breakdown of this data is not made publicly available due to privacy reasons. The Wikimedia Foundation discards raw traffic data after a short retention window, but it collects and retains aggregate historical pageview counts with a geographic breakdown, dating back to 2015 (https://wikitech.wikimedia.org/wiki/Analytics/Data_Lake/Traffic/Pageview_hourly). The pageview data with geographical aggregations used in this study provides the total view counts and the following information for each page: hour, day, year, city, subdivision, country, access method (desktop, mobile or mobile app). Access to this nonpublic pageview data was granted from the Wikimedia Foundation under a non-disclosure agreement as part of its formal collaboration policy. For the analysis conducted in this study, we selected the pageview counts that were localized in the United States only, from January 1, 2016 until December 31, 2016, and aggregated on a weekly basis.

#### Web news

Data were downloaded from the Global Database of Events, Language and Tone (GDELT - http://www.gdeltproject.org), available on Google Cloud Platform. The GDELT is created from real-time translation of worldwide news into 65 languages and updated every 15 minutes. Whenever GDELT detects a news report breaking anywhere the world, the report is then translated, processed to identify all events, counts, quotes, people, organizations, locations, themes, emotions, relevant imagery, video, and embedded social media posts. All the information is made available through an API. In our study, we collected through the Google Cloud Platform all news items written in English and published online in 2016, which mentioned the words “Zika” and “United States” together. The complete query is provided in the Supplementary Information. The dataset contains a total of 112,706 news items from 7,737 different Web news outlets. Metadata associated to each news item allow to select only news mentioning a specific geographic entity beyond the United States, such as states, counties or cities.

#### TV captions

Data were downloaded from the the TV News Archive (https://archive.org/details/tv which is a research library service launched in September 2012. The service is provided by the Internet Archive which, among other sources, collects and preserves television news. The TV News Archive repurposes closed captioning to enable users to search, quote and borrow U.S. TV news programs. For this study, we collected TV news items by searching all mentions of the word “Zika” in the closed captions of any TV News show aired in the United States in 2016, available from the Archive. For each item, the following information is provided: time, TV station, TV program, text snippet of the caption. In total, the dataset comprises 23,855 timestamped mentions of the word “Zika” from 1,410 different TV programs, both in English and Spanish, aired by 64 U.S. TV stations.

#### Zika case notification data

Incidence data of the Zika virus in the United States was collected from the weekly reports published by the Center for Disease Control (CDC). The reports and the associated data were made publicly available by the CDC on GitHub (https://github.com/cdcepi/zika). The CDC epidemiological reports provide the cumulative number of Zika cases by State, starting from February 24, 2016. Additional case counts of January-February 2016 were extracted from CDC official media releases and included in the dataset.

### Model

We model the weekly number of pageview counts to Zika-related Wikipedia pages in each state with a linear regression of the form

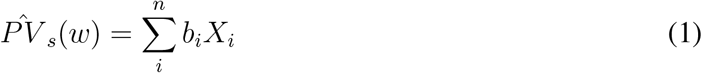

where 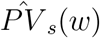 is the Wikipedia page view count in state *s* on week *w*, rescaled by the state population. The rescaling of pageview data takes the form:

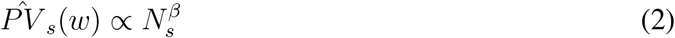

where *N*_*s*_ is the state population and *β* = 1.1397 is a scaling exponent independently estimated on the total volume of pageviews in each state by adopting the probabilistic framework of Leitão et al.^47^. By K-fold (*k* = 10) and leave-one-out cross validation, we test the performance of the model considering different linear combinations of features *X*_*i*_. Specifically, we considered as model features the weekly media timelines *Y*(*w*), where *Y* = TV, Web or Web_*state*_, and Web_*state*_ represents the selection of Web news mentioning only a specific state name together with the word “Zika”. To take into account the saturation effect due to media exposure, we also considered an exponentially decaying function of the media timelines *Y* (*Y* = TV, Web):

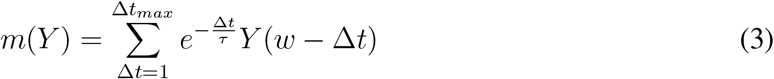

where *τ* is a free parameter, setting the memory time scale, and Δ*t*_*max*_ is defined by the total length of the time series up to week *w* (Δ*t*_*max*_ = *w*). Thus, the full model with all the 5 features under consideration takes the following form:

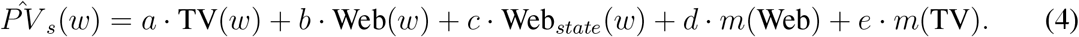

## Acknowledgements

We gratefully acknowledge the Wikimedia Foundation for supporting this work through their formal collaboration and open access policies.

## Contributions

MT, AP, DP and CC designed the study. MT, AP contributed to data collection. MT, AP and CC contributed to data analysis. MT and CC drafted the manuscript. All authors reviewed and approved the final version of this manuscript and take accountability for all aspects of the manuscript.

## Correspondence

Correspondence and requests for materials should be addressed to CC

## Data availability

Anonymized data will be made available upon publication.

## References

1. Jacobs, W., Amuta, A. O. & Jeon, K. C. Health information seeking in the digital age: An analysis of health information seeking behavior among us adults. Cogent Social Sciences 3, 1302785 (2017).

2. Fox, S. & Duggan, M. Health online 2013. Washington, DC: Pew Internet & American Life Project (2013).

3. Ferguson, N. Capturing human behaviour. Nature 446, 733–733 (2007).

4. Poletti, P., Caprile, B., Ajelli, M., Pugliese, A. & Merler, S. Spontaneous behavioural changes in response to epidemics. Journal of theoretical biology 260, 31–40 (2009).

5. Funk, Sebastian and Salathé, Marcel and Jansen, Vincent. Modelling the influence of human behaviour on the spread of infectious diseases: a review. Journal of The Royal Society Interface 7, 1247–1256 (2010).

6. Bauch, C. T. & Galvani, A. P. Social factors in epidemiology. Science 342, 47–49 (2013).

7. Perra, Nicola and Balcan, Duygu and Gonçalves, Bruno and Vespignani, Alessandro. Towards a characterization of behavior-disease models. PLoS One 6, e23084 (2011).

8. Funk, S. et al. Nine challenges in incorporating the dynamics of behaviour in infectious diseases models. Epidemics 10, 21–25 (2015).

9. Lessler, J. et al. Assessing the global threat from zika virus. Science 353, aaf8160 (2016).

10. Mlakar, J. et al. Zika virus associated with microcephaly. N Engl J Med 2016, 951–958 (2016).

11. World Health Organization. WHO Director-General summarizes the outcome of the Emergency Committee regarding clusters of microcephaly and Guillain-Barré syndrome (2016). URL http://www.who.int/mediacentre/news/statements/2016/emergency-committee-zika-microcephaly/en/.

12. World Health Organization. Fifth meeting of the Emergency Committee under the International Health Regulations (2005) regarding microcephaly, other neurological disorders and Zika virus (2016). URL http://www.who.int/mediacentre/news/statements/2016/zika-fifth-ec/en/.

13. Malone, R. W. et al. Zika virus: medical countermeasure development challenges. PLoS Negl Trop Dis 10, e0004530 (2016).

14. Ferguson, N. M. et al. Countering the Zika epidemic in Latin America. Science 353, 353–354 (2016).

15. Zhang, Q. et al. Spread of zika virus in the americas. Proceedings of the National Academy of Sciences 114, E4334–E4343 (2017).

16. Harvard T.H. Chan School of Public Health. Many U.S. families considering pregnancy dont know Zika facts (2016). URL https://www.hsph.harvard.edu/news/press-releases/zika-virus-awareness-pregnant-women/.

17. Shih, T.-J., Wijaya, R. & Brossard, D. Media coverage of public health epidemics: Linking framing and issue attention cycle toward an integrated theory of print news coverage of epidemics. Mass Communication & Society 11, 141–160 (2008).

18. Young, M. E., Norman, G. R. & Humphreys, K. R. Medicine in the popular press: the influence of the media on perceptions of disease. PLoS One 3, e3552 (2008).

19. Collinson, S. & Heffernan, J. M. Modelling the effects of media during an influenza epidemic. BMC public health 14, 376 (2014).

20. Collinson, S., Khan, K. & Heffernan, J. M. The effects of media reports on disease spread and important public health measurements. PloS one 10, e0141423 (2015).

21. Mitchell, L. & Ross, J. V. A data-driven model for influenza transmission incorporating media effects. Open Science 3, 160481 (2016).

22. García-Gavilanes, R., Mollgaard, A., Tsvetkova, M. & Yasseri, T. The memory remains: Understanding collective memory in the digital age. Science Advances 3, e1602368 (2017).

23. Ferron, M. & Massa, P. Beyond the encyclopedia: Collective memories in wikipedia. Memory Studies 7, 22–45 (2014).

24. Shacham, E., Nelson, E. J., Hoft, D. F., Schootman, M. & Garza, A. Potential high-risk areas for zika virus transmission in the contiguous united states. American journal of public health 107, 724–731 (2017).

25. United States Census Bureau. Annual Estimates of the Resident Population for the United States, Regions, States, and Puerto Rico: April 1, 2010 to July 1, 2016. URL https://www2.census.gov/programs-surveys/popest/tables/2010-2016/state/totals/nst-est2016-01.xlsx.

26. Funk, Sebastian and Gilad, Erez and Wtkins, Chris and Jansen, Vincent A.A. The spread of awareness and its impact of epidemic outbreaks. Proceeding of the National Academy of Science 106, 6872–6877 (2009).

27. King, G., Schneer, B. & White, A. How the news media activate public expression and influence national agendas. Science 358, 776–780 (2017).

28. Young, M. E., King, N., Harper, S. & Humphreys, K. R. The influence of popular media on perceptions of personal and population risk in possible disease outbreaks. Health, risk & society 15, 103–114(2013).

29. Towers, S. et al. Mass media and the contagion of fear: the case of ebola in america. PloS one 10, e0129179 (2015).

30. Poletto, C., Boëlle, P.-Y. & Colizza, V. Risk of mers importation and onward transmission: a systematic review and analysis of cases reported to who. BMC infectious diseases 16, 448 (2016).

31. Smith, K. C. et al. Understanding newsworthiness of an emerging pandemic: International newspaper coverage of the h1n1 outbreak. Influenza and other respiratory viruses 7, 847–853 (2013).

32. GeiB, S., Leidecker, M. & Roessing, T. The interplay between media-for-monitoring and media-for-searching: How news media trigger searches and edits in wikipedia. New Media & Society 18, 2740–2759 (2016).

33. García-Gavilanes, R., Tsvetkova, M. & Yasseri, T. Dynamics and biases of online attention: the case of aircraft crashes. Royal Society open science 3, 160460 (2016).

34. Bragazzi, N. L. et al. Global reaction to the recent outbreaks of zika virus: Insights from a big data analysis. PloS one 12, e0185263 (2017).

35. Alicino, C. et al. Assessing ebola-related web search behaviour: insights and implications from an analytical study of google trends-based query volumes. Infectious diseases of poverty 4, 54 (2015).

36. Salathe, M. et al. Digital epidemiology. PLoS computational biology 8, e1002616 (2012).

37. Althouse, B. M. et al. Enhancing disease surveillance with novel data streams: challenges and opportunities. EPJ Data Science 4, 17 (2015).

38. McIver, D. & Brownstein, J. Wikipedia usage estimates prevalence of influenza-like illness in the United States in near real-time. Plos Computational Biology 10, e1003581 (2014).

39. Generous, N., Fairchild, G., Deshpande, A., Del Valle, S. Y. & Priedhorsky, R. Global disease monitoring and forecasting with wikipedia. PLoS computational biology 10, e1003892 (2014).

40. Priedhorsky, R. et al. Measuring global disease with wikipedia: Success, failure, and a research agenda. In Proceedings of the 2017 ACM Conference on Computer Supported Cooperative Work and Social Computing, 1812–1834 (ACM, 2017).

41. Keegan, B., Gergle, D. & Contractor, N. Hot off the wiki: dynamics, practices, and structures in Wikipedia’s coverage of the tohoku catastrophes. International Symposium on Wikis 105–113 (2011).

42. Ratkiewicz, J., Fortunato, S., Flammini, A., Menczer, F. & Vespignani, A. Characterizing and modeling the dynamics of online popularity. Physical Review Letters 105 (2010).

43. Osborne, M., Petrović, S., McCreadie, R., Macdonald, C. & Ounis, I. Bieber no more: First story detection using Twitter and Wikipedia. Proceedings of the Workshop on Time-aware Information Access (2012).

44. Georgescu, M., Kanhabua, N., Krause, D., Nejdl, W. & Siersdorfer, S. Extracting event-related information from article updates in wikipedia. In European Conference on Information Retrieval, 254–266 (Springer, 2013).

45. Tausczik, Y., Faasse, K., Pennebaker, J. W. & Petrie, K. J. Public anxiety and information seeking following the H1N1 outbreak: blogs, newspaper articles, and Wikipedia visits. Health communication 27, 179–185 (2012).

46. Springborn, M., Chowell, G., MacLachlan, M. & Fenichel, E. P. Accounting for behavioral responses during a flu epidemic using home television viewing. BMC Infectious Diseases 15, 21 (2015).

47. Leitao, J. C., Miotto, J. M., Gerlach, M. & Altmann, E. G. Is this scaling nonlinear? Royal Society open science 3, 150649 (2016).

